# Variation in hypoxia tolerance is driven by evolutionary changes in gene regulation in the eastern oyster

**DOI:** 10.64898/2026.10.28.762489

**Authors:** Katherine M. Eaton, Rujuta V. Vaidya, Jerome La Peyre, Jennifer B. Pollack, Kyle A. Sirovy, Morgan W. Kelly

**Affiliations:** Department of Biological Sciences, Louisiana State University, Baton Rouge, LA, 70803, USA; School of Animal Sciences, Louisiana State University Agricultural Center, Baton Rouge, LA, 70803, USA; Harte Research Institute, Texas A&M University-Corpus Christi, Corpus Christi, TX, 78412, USA

**Keywords:** population genomics, low-coverage whole-genome sequencing, climate change, stress tolerance, transcriptomics

## Abstract

Understanding the genomic basis of stress tolerance is crucial for predicting biological responses to environmental change. Here, we investigated the genomic basis of divergence in hypoxia tolerance between two populations of eastern oyster (*Crassostrea virginica*) from Louisiana and Texas, USA. We combined low-coverage whole-genome sequencing data with previously generated transcriptomic evidence to understand the mechanisms underlying population-specific variation in hypoxia tolerance. Genome-wide analyses revealed substantial population differentiation (global F_ST_ = 0.21), indicative of a clear genetic break between our study populations. We combined estimates of genomic differentiation (F_ST_) with diversity metrics to generate a set of exploratory candidate regions showing signatures consistent with recent selection. Several candidate genes identified through this approach were associated with the maintenance of homeostasis under environmental stress. We additionally tested whether genes previously identified as differentially expressed (DE) under hypoxia exhibit elevated genomic divergence between populations. While these hypoxia DEGs did show higher than expected between-population differentiation, this pattern also holds for other environmentally responsive genes. This suggests that transcriptomically responsive genes in general may generally show elevated divergence among populations, possibly due to a weaker effect of purifying selection. Furthermore, we found that hypoxia DEGs did not show an enrichment of nonsynonymous fixed differences (p = 0.9155, as compared to the genome-wide background), suggesting that the elevated divergence seen in these genes is not driven by rapid protein-coding evolution. Together, our results support a model in which population-scale differences in hypoxia tolerance occur primarily through evolution of gene regulation acting on environmentally responsive pathways.

## Introduction

Under anthropogenic climate change, coastal ecosystems are becoming increasingly threatened by hypoxic events, where dissolved oxygen concentrations dip below the threshold required to sustain aerobic marine life (Diaz and Rosenberg 2008; Breitburg et al. 2018). Coastal hypoxia arises through a series of interactions among several climate change-driven factors, including warming water temperatures, which reduce oxygen solubility in water (Shaffer et al. 2009; Keeling et al. 2010; Schmidtko et al. 2017; Breitburg et al. 2018). Beyond this reduced oxygen solubility, climate-associated changes in water column stratification (Chan et al. 2008) and increases in eutrophication (Diaz and Rosenberg 2008) further exacerbate hypoxia, leading to conditions that may severely constrain aerobic metabolism. As dissolved oxygen declines, marine organisms can experience disruptions to growth, reproduction, behavior, and survival, making hypoxia an increasingly important selective pressure in nearshore environments (Taylor and Spicer 1988; Breitburg 2002; Gray et al. 2002; Pollock et al. 2007). Understanding how populations respond and adapt to recurring hypoxic stress is therefore critical for predicting biological responses to ongoing environmental change.

Natural variation in abiotic stress tolerance among populations provides an excellent opportunity to investigate the genomic mechanisms underlying adaptation to environmental change. Populations of the same species inhabiting distinct environments may evolve divergent phenotypes through selection acting on protein-coding sequences, regulatory regions, or combinations of both (Witkopp and Kalay 2012; Savolainen et al. 2013; Ahrens et al. 2019; Kelly 2019; Hao et al. 2023). The relative contribution of regulatory versus coding changes towards local adaptation depends on a variety of factors, including the genetic architecture and biological complexity of the trait in question, as well as the environmental context (Jones et al. 2012; Yeaman 2022). Physiological responses to environmental stress may be particularly likely to involve regulatory changes because they often require coordinated transcriptomic shifts across many genes and pathways (Jones et al. 2012). However, distinguishing the extent to which regulatory and coding changes play a role in adaptive divergence remains a central question in evolutionary genomics, which can be addressed through comparative analysis of populations that show natural variation in stress tolerance.

The eastern oyster, *Crassostrea virginica*, presents a valuable system for investigating the genomic basis of adaptation to hypoxia stress, as populations across its range experience substantial environmental heterogeneity and show natural variation in their response to changing oxygen availability (Byers et al. 2015; Furr et al. 2021; Coxe et al. 2023; Crable et al. 2024). Notably, Coxe et al. (2023) and Crable et al. (2024) leveraged this natural population variation in a common-garden experiment to characterize transcriptional responses to hypoxia in oysters originating from two distinct localities (Upper Laguna Madre, Texas, and Calcasieu Lake, Louisiana) that show variation in abiotic conditions. They found population-specific mortality rates under hypoxia, with oysters originating from Louisiana showing greater sensitivity to low oxygen than their Texas counterparts (Coxe et al. 2023; Crable et al. 2024). Furthermore, they identified several thousand genes that were differentially expressed (DE) in response to hypoxia, with many of these genes showing population-specific expression patterns (e.g., they were DE in one population under hypoxia, but not the other). The common-garden design of this study provides strong evidence that the population-specific transcriptional responses of these genes are associated with underlying population genetic differences. These experimentally identified hypoxia-responsive genes therefore provide a biologically informed set of candidate loci for investigating the genomic basis of divergence in hypoxia tolerance. Because these genes are known to respond to hypoxic stress, we predict that they may also show elevated genomic divergence between populations relative to the genomic background.

In the present study, we combined low-coverage whole-genome sequencing with the transcriptomic data generated by Crable et al. (2024) to investigate the genomic basis of population divergence in hypoxia tolerance between populations of *C. virginica* from Upper Laguna Madre, Texas and Calcasieu Lake, Louisiana (Fig. 1A). We characterized genome-wide population structure and identified regions of the genome showing elevated between-population differentiation using traditional outlier scans. We then tested whether genes previously identified as hypoxia responsive by Crable et al. (2024) showed elevated divergence relative to the genomic background, with particular emphasis on genes that showed population-specific transcriptional responses to hypoxia. Finally, we evaluated whether candidate hypoxia-responsive genes showed evidence of elevated rates of protein-coding evolution, to determine whether divergence was primarily driven by protein-coding or regulatory changes. Taken together, these analyses provide insight into the genomic mechanisms underlying population differences in hypoxia tolerance and demonstrate the value of integrating transcriptomic and population genomic tools towards a mechanistic understanding of stress tolerance evolution.

**Fig. 1.**
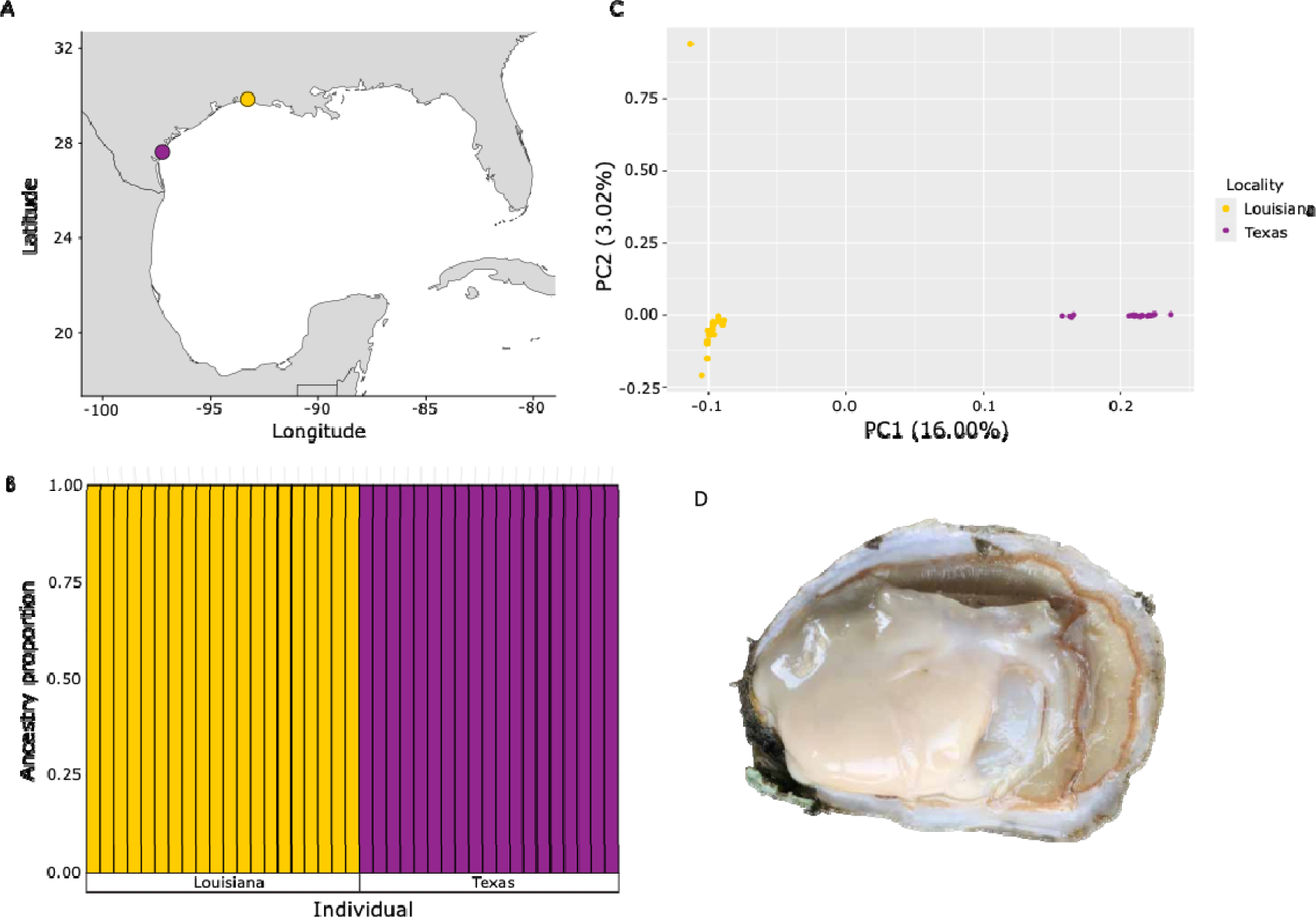
Genome-wide patterns of differentiation among two oyster populations. A. Map of sampling locations for the two oyster populations used in this study (yellow – Calcasieu Lake, Louisiana; purple – Upper Laguna Madre, Texas). B. NGSAdmix plot showing the estimated ancestry proportion for each individual in the study. Individuals are grouped by their location of origin. C. PCA of linkage-pruned SNPs as generated by PCAngsd. Individuals are colored by their location of origin. D. Image of an adult *Crassostrea virginica*

## Methods

### Sample collection

We collected a total of 40 adult eastern oysters (*Crassostrea virginica*) from two populations that show natural variation in their response to hypoxia stress. Individual oysters were collected from Calcasieu Lake, Louisiana (n = 20, 29°50′58.02′′ N, 93°17′1.32′′ W) on January 12, 2016 and from Upper Laguna Madre, Texas (n = 20, 27°37′38′′ N, 97°13′59′′ W) on September 15, 2017 (Fig. 1A). Following collections, oysters were transported live to Louisiana State University (LSU) or Texas A&M University at Corpus Christi where oyster gill tissue was dissected, immersed in 95% ethanol, and stored at −80°C until DNA extraction.

### DNA extraction and sequencing

DNA was extracted from gill tissue using Omega EZNA Tissue DNA kits (Omega Bio-Tek), following the manufacturer’s instructions. The yield and quantity were assessed using a NanoDrop 2000 spectrophotometer and Agilent Bioanalyzer provided by the Genomics Facility at LSU. We used sonication to fragment the DNA and prepared a separate, duel-indexed library for each sample using NEBNext library prep kits for Illumina. The libraries were sent to Novogene (Beijing, China) for low coverage whole-genome sequencing on the Illumina HiSeq X platform, with 150-bp paired-end reads.

### Read mapping

Raw reads were quality-checked using FastQC v. 0.12.1 (Andrews 2010), and adapters were trimmed with Trimmomatic v. 0.39 (Bolger et al. 2014), with the following options: ILLUMINACLIP:TruSeq3-PE-2.fa:2:30:10 LEADING:3 TRAILING:3 SLIDINGWINDOW:4:15 MINLEN:40. Trimmed reads were re-assessed for quality control, again with FastQC (Andrews 2010). We mapped trimmed reads to the *C. virginica* reference genome (v. C_virginica-3.0, NCBI Accession #: GCF_002022765.2) using Bowtie2 v. 2.5.1 (Langmead and Salzberg 2012) with the --very-sensitive option. Alignment files were converted to BAM format, filtered, and sorted with samtools v. 1.19 (Li et al. 2009; Danecek et al. 2021), removing unmapped reads (-F 4), and reads with a mapping quality score < 20 (-q 20). PCR duplicates were removed using Picard MarkDuplicates v. 3.1.1 (Picard Toolkit 2019), and overlapping paired-end reads were clipped using bamUtil clipOverlap v. 1.0.15 (Jun et al. 2015).

### Genotype likelihood estimation and analysis of genome-wide population structure

We estimated genotype likelihoods (GLs) from the final BAM files using ANGSD v. 0.941-26-g6b5d906 with htslib 1.22.1-2-g199e45c6, using the following options: -uniqueOnly 1 -remove_bads 1 -only_proper_pairs 1 -trim 0 -C 50 -minMapQ 20 -minQ 20 -minInd 5 - setMinDepthInd 1 -setMinDepth 5 -doCounts 1 -doCheck 0 -GL 1 -doMajorMinor 1 -doGlf 2 (Korneliussen et al. 2014). We then filtered this GL file to call bi-allelic SNPs, also using ANGSD. For SNP calling, only sites at which at least 20 individuals had ≥1 read covering that site were retained, and any sites with < 20 reads across all samples were discarded (filters: - minInd 20 -setMinDepthInd 1 -setMinDepth 20). We set the p-value filter for polymorphic sites to p<1e-6. Then, to generate a dataset of putatively unlinked SNPs for an initial assessment of genome-wide population structure, we extracted the called SNPs from the genotype likelihood file. These data were formatted and used in the program ngsLD v. 1.2.1 (Fox et al. 2019), to evaluate linkage disequilibrium (LD) decay. LD decay results were plotted using a custom R script and used to determine optimal linkage pruning parameters. We then used the program prune_graph v. 0.3.4 (Vieira 2022) to perform linkage pruning, removing SNP pairs separated by <50kb with a pairwise R^2^ value >0.5. This yielded the pruned dataset of SNPs, for analyses of neutral genetic differentiation. We generated a PCA on the pruned dataset using the program PCAngsd v. 1.36.4 (Meisner and Albrechtsen 2018), which was then visualized in R v. 4.4.1. We also assessed individual admixture proportions with NGSadmix (Skotte et al. 2013), testing K values from 1 to 8 with 20 iterations. We used the Evanno method (Evanno et al. 2005) to identify the optimal K value (i.e., number of genetic clusters) as K = 2, and then visualized the results for K = 2 in R v. 4.4.1.

### Analyses of within- and between-population genomic diversity

To assess patterns of population divergence across the genome, we estimated population-specific site allele frequencies (SAFs) and minor allele frequencies (MAFs) using ANGSD, applying per-population depth and individual coverage filters (i.e., -setMinDepth 10 -minInd 10; all other filtering parameters remain the same as above). The 2-D folded site frequency spectrum (SFS) was then calculated from the SAFs using the realSFS utility in ANGSD. Per-site F_ST_ estimates were then calculated from the SAFs and SFS using the Bhatia et al. (2013) F_ST_ estimator (-whichFst 1), also in realSFS. To characterize spatial patterns of differentiation across the genome, F_ST_ was additionally calculated in 10kb sliding windows, with a step size of 1kb, using the command fst stats2 in realSFS.

We estimated within-population nucleotide diversity (π) by generating the 1-D folded SFS for each population, and then using the command realSFS saf2theta to estimate per-site thetas within a population from the population-specific SAF, leveraging the population-specific 1-D SFS as a prior. Window-based summaries of population nucleotide diversity were then calculated in 10kb sliding windows with a step size of 1kb, using the ANGSD utility thetaStat (do_stat command).

### Genome-wide scans for local adaptation

Our initial approach to identifying regions of the genome putatively involved in hypoxia tolerance was to carry out an exploratory genome-wide F_ST_ scan. Here, we plotted windowed F_ST_ estimates (as calculated in section 2.5) by their genomic location on a Manhattan plot in R v. 4.4.1. We classified regions as outliers if their windowed F_ST_ was in the top 1% of genome-wide F_ST_ values (F_ST_ >= 0.759), merging overlapping regions. We then tested whether genes within these outlier regions were significantly enriched for certain functions via a Gene Ontology (GO) enrichment test. Specifically, we used custom code from Wright et al. (2015) to identify GO categories that were significantly overrepresented in our outlier regions, using a 5% false discovery rate (FDR) cutoff.

To better predict genomic regions that have undergone possible recent selection in one of our two populations, we applied π filters to identify areas of the genome that show a localized reduction in diversity in one population, while also showing high between-population divergence (F_ST_). We calculated the log-transformed π ratio between populations as log_2_(πLouisiana / πTexas), and then defined π outlier regions as those where π was at least 4-fold different between populations (corresponding to a log-transformed π ratio of < −2 or > 2). We then used these π outlier thresholds, combined with a slightly relaxed F_ST_ threshold (top 5% of genome-wide values, F_ST_ >= 0.637), to identify regions that show signatures of recent population divergence and possible selection. As above, we merged overlapping regions. While the top 1% F_ST_ threshold used in our initial genome-wide scan represents a commonly applied cutoff for outlier detection, it is inherently arbitrary. By relaxing the F_ST_ threshold to the top 5%, we retained a more inclusive set of highly diverged regions, to which we could then apply the secondary π ratio filter. This approach ensures candidate regions show convergent evidence of both elevated between-population differentiation and a localized reduction of π in one population, consistent with a recent selective sweep. Candidate regions identified using this approach were visualized on a “meadow plot” (as described by Duffin et al. 2025 preprint), where F_ST_ was plotted by genomic location, and points were colored based on their log-transformed π ratio, and will henceforth be referred to as “meadow plot outliers”. We carried out GO enrichment analyses on these meadow plot outliers by separating outlier regions into those with depressed π in Louisiana and those with depressed π in Texas and testing those sets separately.

### Transcriptomic prior approach to detecting regions putatively under selection

To complement the genome-wide scans described above, we also tested whether genes with known roles in the *C. virginica* hypoxia response show evidence of elevated genomic divergence between populations. To generate our set of candidate hypoxia-responsive genes, we re-analyzed gene expression data originally presented by Crable et al. (2024). Briefly, Crable et al. (2024) reared oysters from the same sampling localities used in the present study (i.e., Calcasieu Lake, Louisiana and Upper Laguna Madre, Texas) under common garden conditions. They then exposed individuals to either control (normoxic) or hypoxic conditions, and took samples to assess gene expression at three time points (72, 96, and 120 hours post-onset of hypoxia, henceforth referred to as time points 1, 2, and 3, respectively). Using the raw count matrix generated by Crable et al. (2024) as input, we ran DESeq2 (Love et al. 2014) independently for each population by time point comparison, yielding lists of genes that were significantly differentially expressed (DE) between hypoxic and normoxic conditions for each of six groups (Louisiana time points 1-3, Texas time points 1-3).

We defined two candidate gene sets with increasing stringency. Our first set of candidate genes (hereafter “hypoxia-responsive”) consisted of all genes that were significantly DE between hypoxic and normoxic conditions in at least one population for at least one time point. These genes represent any gene showing a transcriptional response to hypoxia, regardless of population. The second, more stringent set (hereafter “population-specific hypoxia-responsive”) was restricted to genes that were significantly DE under hypoxia but showed population-specific differences in the magnitude of their expression change. Genes were classified as population-specific hypoxia-responsive candidates if they met either of the following criteria at the same time point: (1) the gene was significantly DE under hypoxia in one population but not the other, or (2) the gene was significantly DE under hypoxia in the same direction in both populations, but the absolute value of the difference in log_2_ fold-change between populations exceeded 1 (i.e., |log_2_FC_Louisiana_ - log_2_FC_Texas_| > 1), indicating a substantial difference in the magnitude of the transcriptional response. This second, more stringent set of genes represents the strongest hypothesized candidates for genomic divergence, as genes that respond differently to hypoxic conditions between populations are most likely to show underlying genetic differences between the tolerant and sensitive populations.

To test whether elevated divergence (if observed) was specific to hypoxia responsive genes or was common to environmentally responsive genes more broadly, we also identified a set of genes that are known to vary in their expression level in response to changes in the environment, but are not necessarily involved in the hypoxia response (henceforth, “environmentally responsive” genes). Theoretically, environmentally responsive genes represent a subset of genes which are more free to vary (in both sequence and expression level) than other, more conserved genes that are under intense purifying selection. If this is the case, we may expect any set of environmentally responsive genes to show elevated divergence relative to the genomic background. We therefore also calculated estimates of between-population genomic divergence for a set of environmentally responsive genes that were significantly DE among individuals of *C. virginica* inhabiting two different field sites along the Gulf Coast of Louisiana, identified by Sirovy et al. (2021). The DEG list discussed here was reproduced from the original dataset analyzed in the R package edgeR (Robinson 2009). This “environmentally responsive” candidate list serves as a negative control, allowing us to confirm whether elevated F_ST_ in our true candidate gene sets is a function of selection on hypoxia tolerance or simply due to elevated divergence in environmentally responsive loci.

To assess whether candidate genes showed significantly elevated F_ST_, we calculated per-gene weighted F_ST_ estimates for each annotated gene in the genome. We calculated weighted F_ST_ by summing the per-site F_ST_ numerator (A) and denominator (B) values across all sites falling within the gene body, plus 10kb upstream and downstream flanking regions, and dividing the summed numerator by the summed denominator. The numerator and denominator values were originally generated in the per-site F_ST_ output generated by realSFS (as described above). This approach yields a single gene-wise F_ST_ estimate for every gene in the genome.

For each candidate gene set, we then tested whether candidate genes showed significantly elevated F_ST_ relative to the genome-wide background of genes using a permutation test. Our test statistic was the difference in mean weighted F_ST_ between candidate genes and all other annotated genes in the genome. To generate a null distribution, we permuted gene labels (i.e., candidate vs. non-candidate) across all annotated genes 10,000 times, each time recalculating the difference in mean F_ST_ between the two groups. The observed test statistic was then compared to this null distribution, and an empirical p-value was calculated as the proportion of permuted differences whose absolute value equaled or exceeded the absolute value of the observed difference. This approach controls for any genome-wide confounders that affect F_ST_ uniformly across genes, as the null distribution is derived from the same set of per-gene F_ST_ values as the observed statistic.

We also tested whether candidate hypoxia-responsive genes showed elevated divergent selection between populations, by estimating the ratio of nonsynonymous to synonymous fixed differences (kN/kS) for candidate and background genes. We identified fixed or nearly-fixed differences between populations using the population-specific MAF files generated by ANGSD (as described above). Sites were filtered to retain only nearly-fixed alleles (minor allele frequency < 0.1) in each population independently, and fixed differences were defined as sites present in both populations where the major allele differed between Louisiana and Texas.

Genome-wide fixed differences were intersected with CDS regions from the *C. virginica* genome annotation (GCF_002022765.2_C_virginica-3.0) using bedtools v. 2.27.1-65-gc2af1e7-dirty (Quinlan 2010). CDS sequences for all longest transcripts were extracted from the reference genome using bedtools getfasta, and multi-exon transcripts were stitched together using a custom python script. Fixed differences between populations were then classified as synonymous or nonsynonymous, again using a custom python script. We then tested whether candidate hypoxia-responsive genes were enriched for nonsynonymous fixed differences relative to background genes using a chi-square test on a 2×2 contingency table of gene category (candidate vs. background) by substitution type (synonymous vs. nonsynonymous). We used this approach rather than calculating a traditional per-gene kN/kS value due to the limited number of fixed differences in coding sequences that were identified between populations.

## Results

### Sequencing

We sequenced 20 individuals each from our two populations, with some variation in sequencing depth by population. Samples from Louisiana had an average of 7,523,811 raw read pairs per individual (± 4,421,059 read pairs, SD), corresponding to a mean of 2.26 Gbp raw sequence data per individual (± 1.33 Gbp, SD). Texas samples had an average of 4,622,132 read pairs per individual (± 1,751,412 read pairs, SD), corresponding to a mean of 1.39 Gbp raw sequence data per individual (± 0.53 Gbp, SD). Following adapter trimming and quality filtering, an average of 7,113,321 read pairs (2.13 Gbp) per individual were retained for the Louisiana samples, and an average of 3,948,467 read pairs (1.18 Gbp) were retained for the Texas samples. Cleaned reads mapped to the *Crassostrea virginica* genome at an average rate of 76.60% for our Louisiana samples (± 0.65%, SD) and 68.85% for our Texas samples (± 2.08%, SD). Following removal of PCR duplicates and final filtering, the average depth for our Louisiana samples was 0.91x, and for our Texas samples was 0.32x. Mean genome coverage for our Louisiana samples was 31.25%, and was 17.85% for our Texas samples. One sample from Texas was identified as an outlier based on its position in a preliminary PCA and was excluded from all further analyses, leaving a final dataset of 39 individuals (20 Louisiana, 19 Texas). All raw sequencing reads have been deposited in NCBI’s Sequence Read Archive (SRA), under BioProject ID PRJNA1510799.

Following GL estimation and SNP filtering, we identified a total of 1,706,770 bi-allelic SNPs, which were then thinned for linkage disequilibrium, resulting in a final putatively unlinked set of 356,816 SNPs, which was used for estimates of population structure.

### Population structure and genomic diversity

PCA revealed clear and strong genomic differentiation between our Louisiana and Texas oyster populations, with no overlap between groups (Fig. 1B). In accordance with the PCA results, NGSAdmix revealed K = 2 as the optimal number of genetic clusters, with all individuals assigned strongly and clearly to their location of origin (Fig. 1C). Together, these results indicate that the Louisiana and Texas oyster populations represent two genetically distinct groups.

The genome-wide global F_ST_ between Louisiana and Texas oyster populations was 0.21, indicating substantial between-population genetic variation. When windowed estimates of F_ST_ were visualized on a Manhattan plot, there was clear genome-wide heterogeneity in differentiation, although nearly every chromosome showed some extremely high F_ST_ peaks (Fig. 2). Within-population estimates of π were similar, with Louisiana oysters showing a mean π of 0.00875, and Texas oysters showing a mean π of 0.00764.

**Fig. 2.**
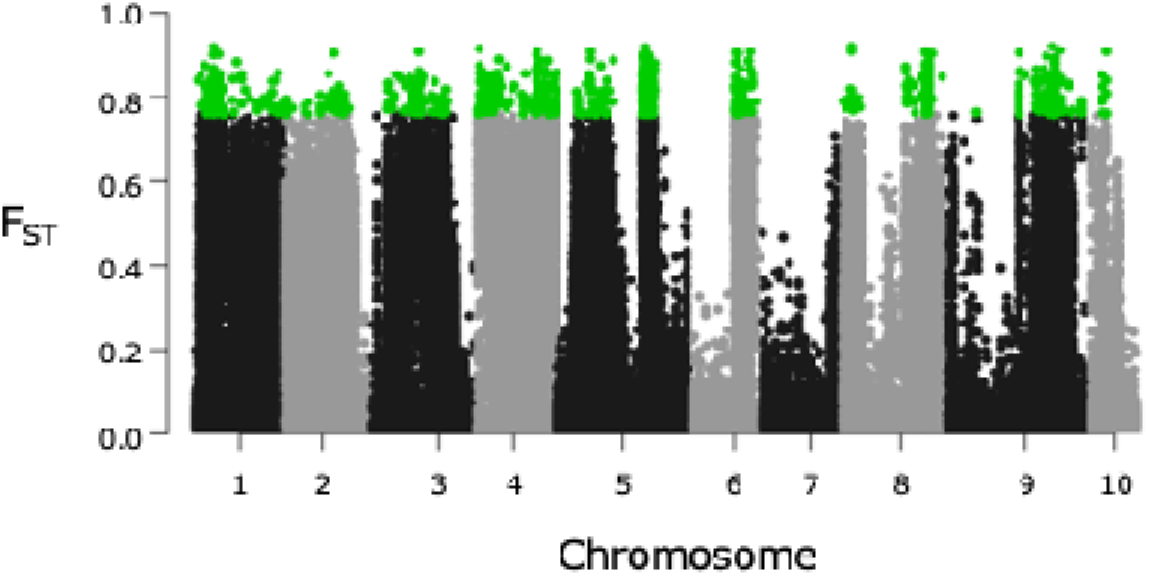
Manhattan plot of genome-wide windowed F_ST_ values between oysters originating from Texas and Louisiana. Outlier windows in the top 1% of genome-wide F_ST_ values are highlighted in green (F_ST_ >= 0.759).

### Genome-wide scans for local adaptation

We identified a total of 547 putative F_ST_ outlier regions distributed across 9 of the 10 chromosomes comprising the *C. virginica* genome (Fig. 2). These outlier regions contained a total of 821 genes (Online Resource 1). GO enrichment analysis of genes within these outlier windows identified 4 significantly enriched GO categories (FDR < 0.05): “guanyl-nucleotide exchange factor activity”, “GTPase regulator activity”, “intracellular signaling cassette”, and “intracellular signal transduction”.

By applying combined F_ST_ and π ratio filters, we generated a meadow plot and detected outlier regions showing elevated genomic divergence and local reductions in nucleotide diversity in one of our two populations (Fig. 3). This approach yielded 50 meadow plot outlier windows, which contained a total of 65 genes (Online Resource 2). Of these 65 genes detected as meadow plot outliers, 47 were also identified as outliers in the top 1% of F_ST_ values. When subdivided based on the directionality of their π ratio bias, 23 of these 50 outlier regions showed depressed in Texas oysters (containing 34 genes, 20 of which overlapped with top 1% of F_ST_ outliers), whereas the remaining 27 showed depressed π in Louisiana oysters (containing 32 genes, 28 of which overlapped with top 1% of F_ST_ outliers). One large gene, *ras guanyl-releasing protein 3-like* (LOC111129997), which was approximately 34 kb in length, overlapped two significant meadow plot outlier regions on chromosome 4 - one of which showed depressed π in Louisiana, and the other of which showed depressed π in Texas.

**Fig. 3.**
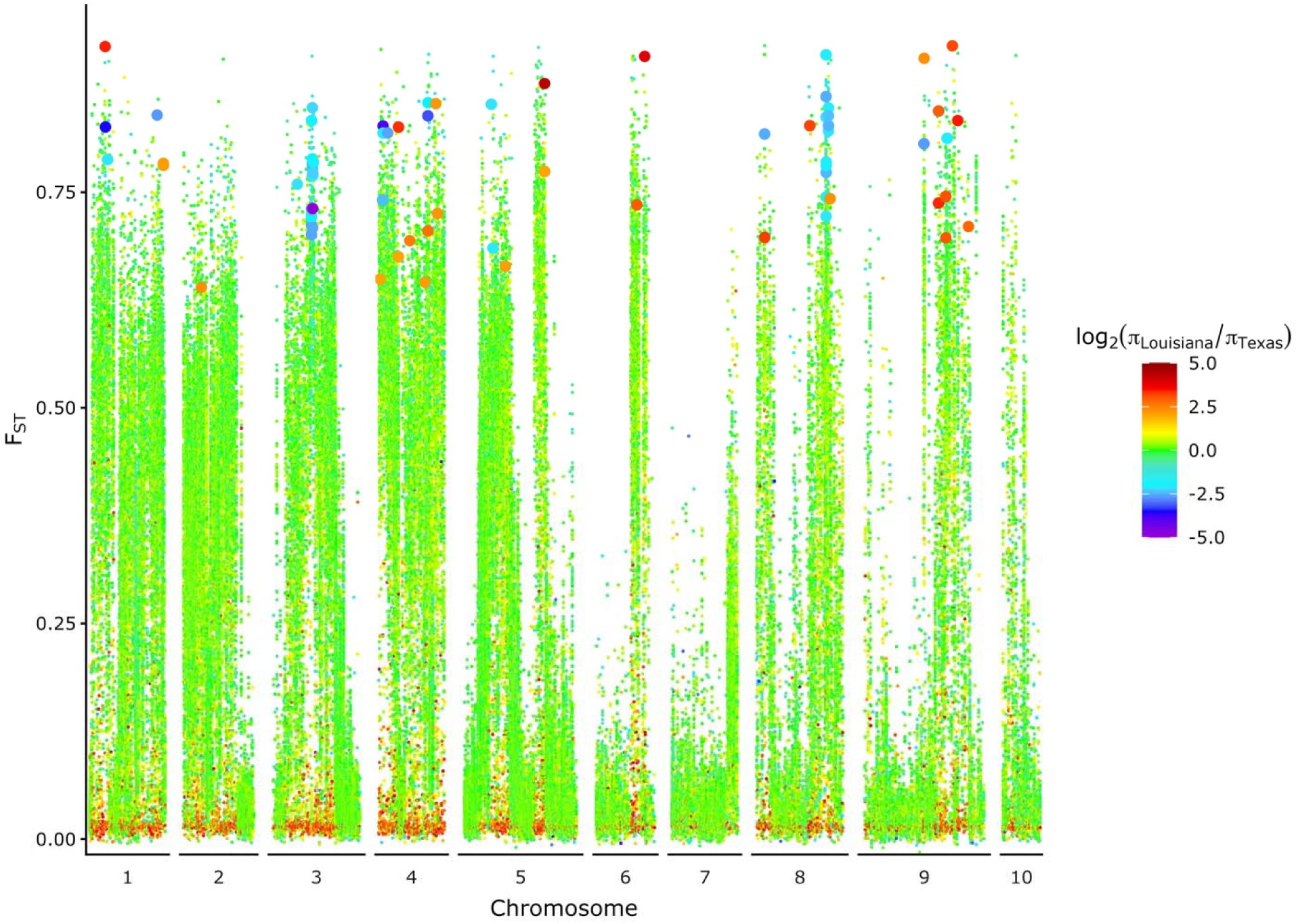
Meadow plot showing windowed genome-wide F_ST_ estimates, colored by their log_2_-transformed π ratio values. Warmer colored points indicate regions with depressed nucleotide diversity in the Texas population, whereas cooler blues and purples indicate regions with depressed nucleotide diversity in the Louisiana population. Genomic windows that fell into the top 5% of genome-wide F_ST_ values and showed a log_2_-transformed π ratio > 2 or < −2 are shown as larger points, to highlight these “meadow plot significant” outlier regions

Of the 34 meadow plot outlier genes showing depressed π in Texas individuals, there were several genes of functional interest, including *mono [ADP-ribose] polymerase PARP16-like* (LOC111128873), *serine/threonine protein kinase Nek 4-like* (LOC111108722), and *ras-related protein Rab-1A-like* (LOC111131278) (Online Resource 2). GO enrichment analysis, however, did not reveal any significantly enriched GO categories within these genes at an FDR <5%.

We also identified 32 meadow plot outlier genes showing depressed π in Louisiana oysters, including *dihydrolipoyl dehydrogenase, mitochondrial-like* (LOC111109254), *Kruppel-like factor 6* (LOC11122603), *pyruvate carboxylase, mitochondrial-like* (LOC111128687), and *beta,beta-carotene 15,15’-dioxygenase-like* (LOC111100534) (Online Resource 2). GO enrichment analysis of these genes resulted in a single enriched GO term: “intracellular signaling cassette”.

### Candidate gene sets investigated for possible local adaptation

Re-analysis of the Crable et al. (2024) gene expression data identified 5,189 hypoxia-responsive genes (i.e., genes which were significantly DE between hypoxic and normoxic conditions in at least one population during at least one time point). Of these 5,189 genes, 4,820 were further classified as population-specific hypoxia-responsive genes, meaning that they showed population-specific differences in the magnitude of their response to hypoxia. A set of broadly environmentally responsive DEGs not necessarily associated with hypoxia tolerance, generated from an unrelated transcriptomic study by Sirovy et al. (2021), serving as a negative control, consisted of 4,926 genes that were DE among oysters inhabiting different environments. Of these, 4,484 hypoxia-responsive genes, 4,143 population-specific hypoxia-responsive genes, and 4,475 broadly environmentally responsive genes had sufficient coverage to calculate weighted F_ST_ estimates and were retained for permutation testing.

Permutation testing revealed that both hypoxia-responsive and population-specific hypoxia-responsive gene sets showed significantly elevated mean weighted F_ST_ relative to the genomic background (Fig. 4). Hypoxia-responsive genes had a mean weighted F_ST_ that was 0.0396 higher than the background genes (p < 0.0001), and population-specific hypoxia-responsive genes showed a similar F_ST_ elevation of 0.0369 above the background genes (p < 0.0001). However, the broadly environmentally responsive gene set also showed a significant elevation in mean weighted F_ST_ relative to background genes (observed difference = 0.0539, p < 0.0001).

**Fig. 4.**
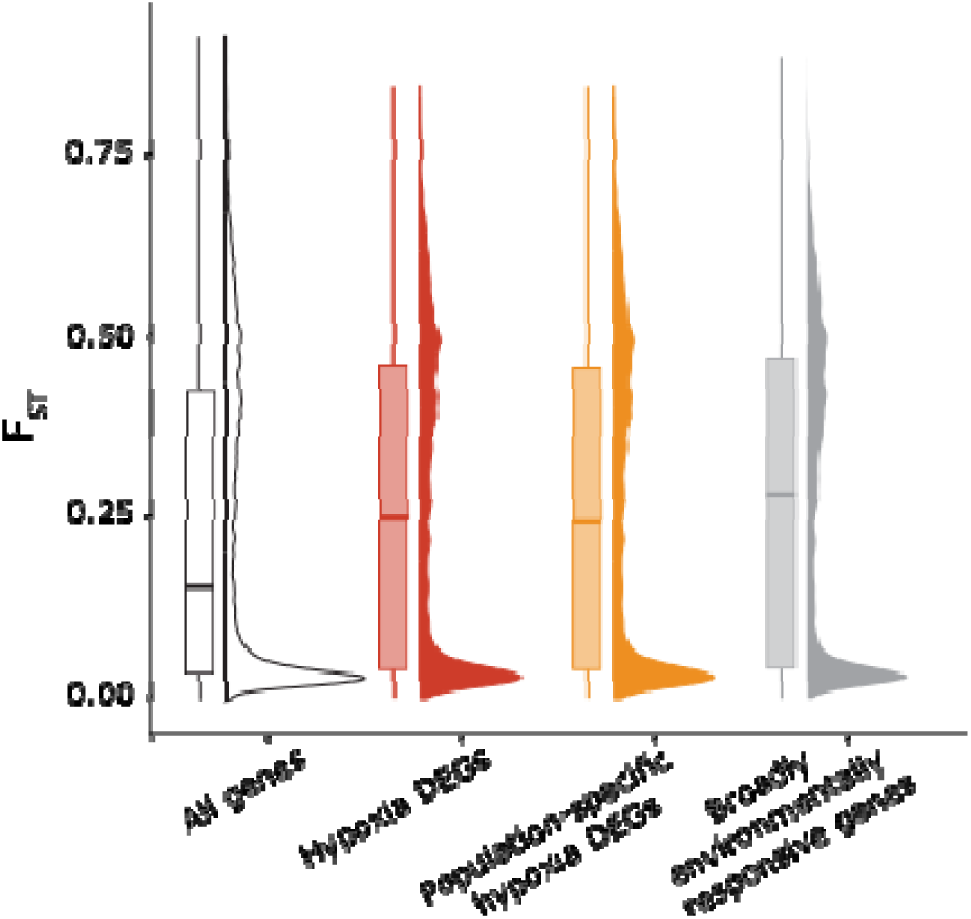
F_ST_ distributions of various candidate gene sets, including all genes (white), significant DEGs under hypoxia (red), population-specific DEGs under hypoxia (orange), and broadly environmentally responsive genes (gray). Hypoxia DEGs, population-specific hypoxia DEGs, and broadly environmentally responsive genes all show significantly higher mean F_ST_ values than the genome-wide average of all genes

We identified a total of 19,336 fixed or nearly-fixed (MAF < 0.1) SNPs between populations, of which 5,726 fell within annotated coding sequences in the *C. virginica* genome. These 5,726 SNPs in coding regions were retained for kN/kS analyses. The number of synonymous and nonsynonymous fixed differences in genes in each of our three candidate gene sets is shown in Online Resource 3. Neither candidate gene set showed a significant enrichment of nonsynonymous fixed differences relative to background genes (hypoxia responsive: *χ*^2^ = 0.011258, df = 1, p = 0.9155, population-specific hypoxia-responsive: *χ*^2^ = 0.13044, df = 1, p = 0.718), nor did the null gene set (*χ*^2^ = 0.16073, df = 1, p = 0.6885).

## Discussion

We investigated the genomic basis of divergence in hypoxia tolerance between eastern oyster populations using new low-coverage whole-genome sequencing data. We found strong genome-wide differentiation between oyster populations from Louisiana and Texas, indicating substantial genetic structure across the Gulf Coast. Against this background of broad genomic differentiation, traditional genomic outlier scans identified hundreds of peaks of genomic divergence, making it difficult to distinguish loci associated with adaptive divergence from those reflecting broader population history. By additionally applying π filters to our F_ST_ outliers, we narrowed our candidate region list into those showing signatures more consistent with recent selection (i.e., high differentiation and a localized reduction in diversity in one population).

Beyond these exploratory analyses, we also found that genes with a known role in the hypoxia response (as previously identified in a transcriptomic study by Crable et al. (2024)) show elevated genomic divergence, as compared to background genes, and that this divergence is not due to increases in rates of protein-coding evolution. Together, these findings suggest that adaptive divergence in hypoxia tolerance may occur primarily through changes in gene regulation acting on environmentally responsive pathways rather than through widespread modification of coding sequence.

### Major genome-wide differentiation points towards a genetic break along the Gulf Coast, consistent with previous work

The Louisiana and Texas oyster populations exhibited substantial genome-wide differentiation (global F_ST_ = 0.21, Fig. 1B,C). This indicates extensive evolutionary divergence between populations, despite the geographic proximity of the two sites (separated by < 500 km, Fig. 1A). This level of differentiation suggests that population history and restricted gene flow have generated broad structure across the genome, consistent with previous work describing high population subdivision in *Crassostrea virginica*, with a notable genetic break along the Gulf Coast of Texas (Anderson et al. 2014). While such divergence creates ample opportunity for local adaptation, it also complicates the interpretation of traditional F_ST_-based outlier scans, which assume that loci under positive selection will show elevated divergence against a relatively homogeneous genetic background (Akey et al. 2002). When baseline differentiation is already high, large numbers of highly differentiated regions make it difficult to distinguish putatively adaptive differences from those that have arisen through neutral processes. Indeed, our exploratory F_ST_-based analysis identified hundreds of high F_ST_ regions across the *C. virginica* genome (Fig. 2), showing that between-population differentiation is widespread, rather than concentrated in a few putatively adaptive regions.

### Nucleotide diversity filters improve traditional outlier scans, yielding several candidate regions under putative selection in hypoxia-tolerant oysters

To improve our detection power of putatively adaptive regions under this highly differentiated background, we combined π and F_ST_ filters to identify genomic regions that showed both high between-population differentiation and a localized reduction in nucleotide diversity in one of our two populations, suggesting a possible recent selective sweep. This additional filtering approach yielded a much smaller set of possible candidate regions more likely to reflect recent evolutionary change than historical divergence. Combining diversity-based metrics like π with differentiation scans has been shown to improve identification of candidate loci under selection in a number of other species, including domestic sheep (Li et al. 2020), pigs (Zhang et al. 2020; Chen et al. 2022), and sea stars (Duffin et al. 2025 preprint).

Using this combined “meadow-plot” style approach, we identified 34 candidate genes under putative selection in hypoxia-tolerant Texas oysters (Fig. 3). Notably, several of these genes are associated with maintenance of cellular homeostasis under stress, including *PARP-16-like* (LOC111128873), *NEK-4-like* (LOC111108722) and *RAB-1A-like* (LOC111131278).

Cellular responses to hypoxia in aquatic organisms frequently involve activation of pathways that mitigate protein misfolding, oxidative damage from free radical accumulation, and disruption of metabolism from reduced oxygen availability (Wu 2002; Ivanina et al. 2016; Ivanina and Sokolova 2016; Montúfar-Romero et al. 2024).

*PARP-16* plays a key role in the endoplasmic reticulum (ER) stress responses by regulating the unfolded protein response (UPR) signaling pathway (Jwa and Chang 2012). The UPR is activated when unfolded proteins accumulate in the ER, a common effect of exposure to hypoxia (Cao and Kaufman 2012). The locus in the meadow plot outlier region containing *PARP-16* that showed the highest between-population differentiation (F_ST_ = 0.82) occurred approximately 5kb upstream of the start of *PARP-16* coding sequence, suggesting a possible change in *PARP-16* regulation in our hypoxia-tolerant Texas oysters.

*NEK-4* has also been implicated in the cellular stress response, playing a role in promoting cell cycle arrest under double-stranded DNA damage (Nguyen et al. 2012). As hypoxia followed by reoxygenation can increase reactive oxygen species production and oxidative damage, including double-stranded DNA breaks, changes in pathways involved in DNA damage tolerance may influence resilience to oxygen limitation (Hadj-Moussa et al. 2022). Beyond its role in genomic maintenance, *NEK-4* has also been linked to mitochondrial physiology, where experimental overexpression of *NEK-4* increases mitochondrial respiration rate and alters cellular metabolic state (Basei et al. 2022). There were three highly differentiated (F_ST_ > 0.90) SNPs that all occurred within a 100 bp region directly upstream of the *NEK-4* start codon, again suggesting that changes in gene regulation may play an important role in between-population variation in hypoxia tolerance.

*RAB-1A* is a regulator of intracellular transport that plays a major role in the stress-induced autophagy pathway, contributing to degradation and recycling of damaged proteins under cellular stress (Gyurkovska et al. 2023). Autophagy is frequently induced under hypoxia and is thought to promote cell survival by maintaining metabolic homeostasis and removing damaged cellular components (Kawsar et al. 2025). Interestingly, *RAB-1A* has undergone gene duplication in the common carp (*Cyprinus carpio*), a species with high tolerance to low oxygen environments (Zhao et al. 2015). Divergence in expression patterns of *RAB-1A* paralogs under hypoxia is thought to contribute to the hypoxia-tolerant phenotype of this fish species (Zhao et al. 2015). Much like *PARP-16* and *NEK-4*, the locus within the meadow plot outlier region containing *RAB-1A* with the highest between-population differentiation (F_ST_ = 0.88) was upstream of the *RAB-1A* start site, again consistent with changes in gene regulation.

### Hypoxia-responsive genes show elevated divergence, but this pattern extends across broadly environmentally responsive loci

To complement the exploratory genome-wide outlier scans described above, we incorporated transcriptomic information from Crable et al. (2024) and tested whether genes known to respond to hypoxic stress exhibit elevated genomic divergence between populations. Consistent with our prediction, genes that were differentially expressed under hypoxic conditions exhibited significantly higher weighted F_ST_ than background genes across the *C. virginica* genome (Fig. 4). This pattern persisted when restricting analyses to genes showing population-specific transcriptional responses under hypoxia (Fig. 4), highlighting that genes involved in population-specific responses to oxygen limitation tend to show signatures of elevated between-population divergence. These findings support our hypothesis that changes in hypoxia-responsive pathways contribute to population differentiation in this species.

Interestingly, we also observed a similar elevation in genomic divergence among our null candidate gene set, consisting of environmentally responsive genes identified in an unrelated transcriptomic study (Fig. 4; Sirovy et al. 2021). This result indicates that elevated between-population divergence is not necessarily unique to hypoxia-responsive genes, instead suggesting that broadly environmentally responsive genes may evolve differently from the genomic background. One possible explanation for this pattern is that genes involved in environmentally induced responses may experience weaker purifying selection than genes required for highly conserved cellular functions. Environmentally responsive genes are expressed conditionally, in specific tissues or environmental contexts, and may therefore tolerate greater variation in both coding sequence and regulation without fatally disrupting organismal performance (Lasky et al. 2014; Kelly 2019; Sirovy et al. 2021). Alternatively, environmentally responsive genes may be more generally subject to spatially divergent selection than other genes in the genome, as selection optimizes expression across varying environments.

Importantly, this interpretation does not preclude hypoxia-responsive genes from contributing to adaptive divergence. Rather, it suggests that environmentally responsive genes may represent a subset of the genome that is generally more evolutionarily flexible, therefore providing particularly accessible substrate for selection to act upon. Under this framework, elevated divergence in hypoxia-responsive genes may reflect both their functional importance in oxygen stress responses and their increased capacity to accumulate regulatory and sequence variation relative to more strongly constrained loci.

### Divergence in hypoxia-responsive genes occurs primarily through regulatory sequence evolution

While both changes in protein-coding sequence and gene regulatory architecture are recognized as important drivers of local adaptation, environmental stress responses are often hypothesized to evolve predominantly through regulatory change because they require coordinated shifts in expression across large networks of interacting genes (Jones et al. 2012; Fraser 2013; Kelly 2019). Despite observing elevated genomic divergence among hypoxia-responsive genes, we found no evidence that these candidate loci exhibited elevated rates of protein-coding evolution. This result suggests that the divergence associated with hypoxia tolerance among our populations occurs primarily outside of coding sequence - in agreement with our exploratory genome scan results that showed high F_ST_ SNPs directly upstream of functionally relevant candidate genes. Cis-regulatory evolution has long been proposed as an efficient method for adaptation because it can alter gene expression in specific tissues or environmental contexts while minimizing pleiotropic effects associated with protein-coding mutations (Wray 2007; Fraser 2013). Such mechanisms may be particularly relevant for hypoxia responses, which involve extensive transcriptional reorganization across metabolic and cellular stress response pathways.

However, our results should not be interpreted as evidence that coding sequence evolution is unimportant for hypoxia adaptation. Due to the nature of our low-coverage data, our kN/kS analysis was necessarily restricted to fixed or nearly fixed differences among populations, and therefore had limited power to detect more subtle patterns of protein evolution, given the relatively small number of fixed SNPs in protein-coding sequence between populations. Future work integrating population genomics with functional approaches, such as chromatin accessibility assays and allele-specific expression analyses will be necessary to confirm whether the regulatory divergence observed here directly contributes to differences in hypoxia tolerance between populations. More broadly, though, our results highlight the value of integrating transcriptomic and population genomic approaches to identify mechanisms of adaptation to environmental stress.

## Conclusion

Our results demonstrate substantial genomic differentiation between populations of *C. virginica* that differ in hypoxia tolerance, and highlight the challenges of identifying adaptive loci in systems with elevated baseline divergence. By integrating population genomic scans with transcriptomic prior information, we identified candidate genomic regions associated with hypoxia-responsive pathways and found that genes involved in the hypoxia response exhibit elevated genomic differentiation relative to background loci. However, the absence of evidence for elevated protein coding divergence among candidate genes suggests that this differentiation is unlikely to be driven primarily by coding sequence evolution. Instead, our findings are most consistent with a model in which adaptive divergence in hypoxia tolerance occurs primarily through changes in gene regulation. Ultimately, this work demonstrates the value of combining population genomic and transcriptomic data to investigate the mechanisms underlying adaptation to climate-driven environmental stress.

## Supporting information

Online Resource 1

Online Resource 2

Online Resource 3

## Compliance with Ethical Standards

This work was funded by a grant from the National Science Foundation (NSF-BioOce 1731710) to Morgan Kelly and Jerome La Peyre. The authors declare no conflict of interest. All applicable international, national, and institutional guidelines for sampling, care, and experimental use of organisms for the study have been followed. No institutional ethical committee review is required for the use of invertebrate animals in research.

## Acknowledgements

This work was funded by the National Science Foundation (NSF-BioOCE 1731710). We are grateful to Abdul Ada and Isabelle Neylan for helpful discussions regarding analyses and results. We thank the Louisiana State University High Performance Computing Facility for access to computing resources used to carry out the analyses included in this study.

## Statements and Declarations

## Author Contributions

All authors contributed to the conceptualization, methodology, and investigation of this project. Formal analysis, validation, data curation, and visualization were performed by Katherine M. Eaton, with advice from Morgan W. Kelly and Rujuta V. Vaidya. Resources were provided by Morgan W. Kelly, Jerome La Peyre, and Jennifer B. Pollack. The original draft was written by Katherine M. Eaton, with assistance from Kyle A. Sirovy and Morgan W. Kelly. All authors contributed to the review and editing of the final manuscript. Project supervision and administration was done by Morgan W. Kelly.

## Data Availability

The genomic sequence data generated during this study has been uploaded to the National Center for Biotechnology Information Sequence Read Archive (NCBI SRA) under BioProject accession number PRJNA1510799. These data are currently embargoed until publication, but will be made available upon article acceptance. Reviewer links can be generated upon request.

## Ethics Approval

All applicable international, national, and institutional guidelines for sampling, care, and experimental use of organisms for the study have been followed. As this study used invertebrate animals, no Institutional Animal Care and Use Committee approval was required by Louisiana State University.

## References

Ahrens CW, Byrne M, Rymer PD (2019) Standing genomic variation within coding and regulatory regions contributes to the adaptive capacity to climate in a foundation tree species. Mol Ecol. 28(10):2502–2516. 10.1111/mec.15092

Akey JM, Zhang G, Zhang K, Jin L, Shriver MD (2002) Interrogating a high-density SNP map for signatures of natural selection. Genome Res. 12:1805–1814. 10.1101/gr.631202

Anderson JD, Karel WJ, Mace CE, Bartram BL, Hare MP (2014) Spatial genetic features of eastern oysters (*Crassostrea virginica* Gmelin) in the Gulf of Mexico: northward movement of a secondary contact zone. Ecol Evol. 4(9):1671–1685. 10.1002/ece3.1064

Andrews S (2010) FastQC: a quality control tool for high throughput sequence data. https://www.bioinformatics.babraham.ac.uk/projects/fastqc/. [Accessed 29 June 2026.]

Basei FL, de Castro Ferezin C, Rodrigues de Oliveira AL, Muñoz JP, Zorzano A, Kobarg J (2022) Nek4 regulates mitochondrial respiration and morphology. FEBS J. 289(11):3262–3279. 10.1111/febs.16343

Bhatia G, Patterson N, Sankararaman S, Price AL (2013) Estimating and interpreting FST: the impact of rare variants. Genome Res. 23(9):1514–1521. 10.1101/gr.154831.113

Bolger AM, Lohse M, Usadel B (2014) Trimmomatic: a flexible trimmer for Illumina sequence data. Bioinformatics. 30(15):2114–2120. 10.1093/bioinformatics/btu170

Breitburg D (2002) Effects of hypoxia, and the balance between hypoxia and enrichment, on coastal fishes and fisheries. Estuaries. 25(4b):767–781. 10.1007/BF02804904

Breitburg D, Levin LA, Oschlies A, et al (2018) Declining oxygen in the global ocean and coastal waters. Science. 359:eaam7240. 10.1126/science.aam7240

Byers JE, Grabowski JH, Piehler MF, Hughes AR, Weiskel HW, Malek J, Kimbro DL (2015) Geographic variation in intertidal oyster reef properties and the influence of tidal prism. Limnol Oceanogr. 60(3):1051–1063. 10.1002/lno.10073

Cao SS, Kaufman RJ (2012) Unfolded protein response. Curr Biol. 22(16):PR622–626. 10.1016/j.cub.2012.07.004

Chan F, Barth JA, Lubchenco J, Kirincich A, Weeks H, Peterson WT, Menge BA (2008) Emergence of anoxia in the California Current Large Marine Ecosystem. Science. 319:920. 10.1126/science.1149016

Chen Y, Wu X, Wang J, et al (2022) Detection of selection signatures in Anqing six-end-white pigs on resequencing data. Genes-Basel. 13(12):2310. 10.3390/genes13122310

Crable EL, Rodriguez H, Vaidya RV, Coxe N, La Peyre JF, Kelly MW (2024) Transcriptomic responses to hypoxia in two populations of eastern oyster with differing tolerance. Front Mar Sci. 11:1462905. 10.3389/fmars.2024.1462905

Coxe N, Casas SM, Marshall DA, La Peyre MK, Kelly MW, La Peyre JF (2023) Differential hypoxia tolerance of eastern oysters from the northern Gulf of Mexico at elevated temperature. J Exp Mar Biol Ecol. 559:151840. 10.1016/j.jembe.2022.151840

Danecek P, Bonfield JK, Liddle J, et al (2021) Twelve years of SAMtools and BCFtools. GigaScience. 10(2):giab008. 10.1093/gigascience/giab008

Diaz RJ, Rosenberg R (2008) Spreading dead zones and consequences for marine ecosystems. Science. 321(5891):926–929. 10.1126/science.1156401

Duffin PJ, Schiebelhut LM, Dawson MN, Wares JP (2025) Genomic separation of Salish Sea and Pacific outer coast populations of the keystone sea star *Pisaster ochraceus*. bioRxiv. 10.1101/2025.07.24.666598

Evanno G, Regnaut S, Goudet J (2005) Detecting the number of clusters of individuals using the software STRUCTURE: a simulation study. Mol Ecol. 14(8):2611–2620. 10.1111/j.1365-294X.2005.02553.x

Fox EA, Wright AE, Fumagalli M, Vieira FG (2019) ngsLD: evaluating linkage disequilibrium using genotype likelihoods. Bioinformatics. 35(19):3855–3856. 10.1093/bioinformatics/btz200

Fraser HB (2013) Gene expression drives local adaptation in humans. Genome Res. 23(7):1089–1096. 10.1101/gr.152710.112

Furr D, Ketchum RN, Phippen BL, Reitzel AM, Ivanina AV (2021) Physiological variation in response to *Vibrio* and hypoxia by aquacultured eastern oysters in the southeastern United States. Integr Comp Biol. 61(5):1715–1729. 10.1093/icb/icab176

Gray JS, Wu RS, Or YY (2002) Effects of hypoxia and organic enrichment on the coastal marine environment. Mar Ecol Progress Series. 238:249–279. 10.3354/meps238249

Gyurkovska V, Murtazina R, Zhao SF, Shikano S, Okamoto Y, Segev N (2023) Dual function of Rab1A in secretion and autophagy: hypervariable domain dependence. Life Sci Alliance. 6(5):e202201810. 10.26508/lsa.202201810

Hadj-Moussa H, Eaton L, Cheng H, Pamenter ME, Storey KB (2022) Naked mole-rats resist the accumulation of hypoxia-induced oxidative damage. Comp Biochem Phys A. 273:111282. 10.1016/j.cbpa.2022.111282

Hao Y, Song G, Zhang YE, Zhai W, Jia C, Ji Y, Tang S, Lv H, Qu Y, Lei F (2023) Divergent contributions of coding and noncoding sequences to initial high-altitude adaptation in passerine birds endemic to the Qinghai-Tibet plateau. Mol Ecol. 32(13):3524–3540. 10.1111/mec.16942

Ivanina AV, Nesmelova I, Leamy L, Sokolov EP, Sokolova IM (2016) Intermittent hypoxia leads to functional reorganization of mitochondria and affects cellular bioenergetics in marine molluscs. J Exp Biol. 219(11):1659–1674. 10.1242/jeb.134700

Ivanina AV, Sokolova IM (2016) Effects of intermittent hypoxia on oxidative stress and protein degradation in molluscan mitochondria. J Exp Biol. 219(23):3794–3802. 10.1242/jeb.146209

Jones FC, Grabherr MG, Chan YF, et al (2012) The genomic basis of adaptive evolution in threespine sticklebacks. Nature. 484:55–61. 10.1038/nature10944

Jun G, Wing MK, Abecasis GR, Kang HM (2015) An efficient and scalable analysis framework for variant extraction and refinement from population scale DNA sequence data. Genome Res. 25(6):918–925. 10.1101/gr.176552.114

Jwa M, Chang P (2012) PARP16 is a tail-anchored endoplasmic reticulum protein required for the PERK- and IRE1*α*-mediated unfolded protein response. Nat Cell Biol. 14:1223–1230. 10.1038/ncb2593

Kawsar MA, Adikari D, Zhang Y (2025) Autophagy in aquatic animals: mechanisms, implications, and future directions. Front Immunol. 16:1612178. 10.3389/fimmu.2025.1612178

Keeling RF, Körtzinger A, Gruber N (2010) Ocean deoxygenation in a warming world. Annu Rev Mar Sci. 2:199–229. 10.1146/annurev.marine.010908.163855

Kelly M (2019) Adaptation to climate change through genetic accommodation and assimilation of plastic phenotypes. Philos T Roy Soc B. 374(1768):20180176. 10.1098/rstb.2018.0176

Korneliussen TS, Albrechtsen A, Nielsen R (2014) ANGSD: analysis of next generation sequencing data. BMC Bioinformatics. 15:356. 10.1186/s12859-014-0356-4

Langmead B, Salzberg S (2012). Fast gapped-read alignment with Bowtie2. Nat Methods. 9(4):357–359. 10.1038/nmeth.1923

Lasky JR, Des Marais DL, Lowry DB, Povolotskaya I, McKay JK, Richards JH, Keitt TH, Juenger TE (2014) Natural variation in abiotic stress responsive gene expression and local adaptation to climate in *Arabidopsis thaliana*. Mol Biol Evol. 31(9):2283–2296. 10.1093/molbev/msu170

Li H, Handsaker B, Wysoker A, Fennell T, Ruan J, Homer N, Marth G, Abecasis G, Durbin R, The 1000 Genome Project Data Processing Subgroup (2009) The Sequence alignment/map (SAM) format and SAMtools. Bioinformatics. 25(16):2078–9. 10.1093/bioinformatics/btp352

Li X, Yang J, Shen M, et al (2020) Whole-genome resequencing of wild and domestic sheep identifies genes associated with morphological and agronomic traits. Nature Commun. 11:2815. 10.1038/s41467-020-16485-1

Meisner J, Albrechtsen A (2018) Inferring population structure and admixture proportions in low-depth NGS data. Genetics. 210(2):719–731. 10.1534/genetics.118.301336

Montúfar-Romero M, Valenzuela-Muñoz V, Valenzuela-Miranda D, Gallardo-Escárate C (2024) Hypoxia in the blue mussel *Mytilus chilensis* induces a transcriptomic shift associated with endoplasmic reticulum stress, metabolism, and immune response. Genes-Basel. 15(6):658. 10.3390/genes15060658

Nguyen CL, Possemato R, Bauerlein EL, Xie A, Scully R, Hahn WC (2012) Nek4 regulates entry into replicative senescence and the response to DNA damage in human fibroblasts. Mol Cell Biol. 32(19):3963–3977. 10.1128/MCB.00436-12

Picard Toolkit (2019) Broad Institute, GitHub Repository. https://broadinstitute.github.io/picard/ [Accessed 29 June 2026.]

Pollock MS, Clarke LMJ, Dubé MG (2007) The effects of hypoxia on fishes: from ecological relevance to physiological effects. Environ Rev. 15:1–14. 10.1139/a06-006

Savolainen O, Lascoux M, Merilä J (2013) Ecological genomics of local adaptation. Nat Rev Genet. 14:807–820. 10.1038/nrg3522

Schmidtko S, Stramma L, Visbeck M (2017) Decline in global oceanic oxygen content during the past five decades. Nature. 547(7641):335–339. 10.1038/nature21399

Shaffer G, Olsen S, Pedersen J (2009) Long-term ocean oxygen depletion in response to carbon dioxide emissions from fossil fuels. Nat Geosci. 2:105–109. 10.1038/ngeo420

Sirovy KA, Johnson KM, Casas SM, La Peyre JF, Kelly MW (2021) Lack of genotype-by-environment interaction suggests limited potential for evolutionary changes in plasticity in the eastern oyster, *Crassostrea virginica*. Mol Ecol. 30:5721–5734. 10.1111/mec.16156

Skotte L, Korneliussen TS, Albrechtsen A (2013) Estimating individual admixture proportions from next generation sequencing data. Genetics. 195(3):693–702. 10.1534/genetics.113.154138

Streisfeld MA, Young WN, Sobel JM (2013) Divergent selection drives genetic differentiation in an R2R3-MYB transcription factor that contributes to incipient speciation in *Mimulus aurantiacus*. PLOS Genet. 9(3):e1003385. 10.1371/journal.pgen.1003385

Taylor AC, Spicer JI (1988) Functional significance of a partial emersion response in the intertidal prawn *Palaemon elegans* (Crustacea: Palaemonidae) during environmental hypoxia. Mar Ecol Prog Ser. 44:141–147.

Vieira FG (2022) prune_graph, GitHub repository. https://github.com/fgvieira/prune_graph [Accessed 29 June 2026.]

Wittkopp PJ, Kalay G (2012) *Cis*-regulatory elements: molecular mechanisms and evolutionary processes underlying divergence. Nat Rev Genet. 13:59–69. 10.1038/nrg3095

Wray G (2007) The evolutionary significance of *cis-*regulatory mutations. Nat Rev Genet. 8:206–216. 10.1038/nrg2063

Wright RM, Aglyamova GV, Meyer E, Matz MV (2015) Gene expression associated with white syndromes in a reef building coral, *Acropora hyacinthus*. BMC Genomics. 16(1):371. 10.1186/s12864-015-1540-2

Wu RSS (2002) Hypoxia: from molecular responses to ecosystem responses. Mar Pollut Bull. 45:35–45. 10.1016/S0025-326X(02)00061-9

Yeaman S (2022) Evolution of polygenic traits under global *vs* local adaptation. Genetics. 220(1):iyab134. 10.1093/genetics/iyab134

Zhang W, Yang M, Zhou M, Wang Y, Wu X, Zhang X, Ding Y, Zhao G, Yin Z, Wang C (2020) Identification of signatures of selection by whole-genome resequencing of a Chinese native pig. Front Genet. 11:566255. 10.3389/fgene.2020.566255

Zhao ZX, Cao DC, Xu J, Xu R, Li JT, Zhang Y, Xu P, Sun XW (2015) Diversification of the duplicated *Rab1a* genes in a hypoxia-tolerant fish, common carp (*Cyprinus carpio*). Comp Biochem Phys B. 188:54–62. 10.1016/j.cbpb.2015.06.007

